# Legacy effects of extreme events

**DOI:** 10.1101/2024.07.09.602699

**Authors:** Danielle Brueggemeier-Singh, Matthijs Vos

## Abstract

Global change is increasing the intensity of environmental extremes. Abiotic extremes include hotter summers and more intense heatwaves^1-2^. Biotic extremes include disproportionate increases in herbivory^3^. Both have well-understood immediate impacts on the plant communities that support life across the globe. However, their longterm effects are difficult to predict, as it is unknown how heat and herbivory differ in the way they create historical legacies. These are changes in a community that still affect dynamics and recovery long after a disturbance has passed. Here we show differences in legacy construction by mild and extreme heatwaves and small- and large-bodied herbivores. We analysed patterns in the population dynamics of a consumer added to replicated primary producer communities after disturbance had ended. We find that the legacy induced by extreme heat drives post-disturbance consumer declines in all replicates. This results in stochastic extinctions as the loss of populations becomes more likely when their density gets close to zero. In contrast, the legacy created by the large-bodied herbivore does not drive extinctions. It diminishes population growth by orders of magnitude across six generations. The observed effects appear after a time-lag that is shorter for extreme heat than for herbivory. Our results show that legacy effects drive complex causal sequences that may involve deterministic and chance components, time-lags, stressor amplification and regime shifts. Insight in how these combine to shape the future is essential for our capacity to accurately forecast and repair the consequences of global change.

---

Reliable predictions are essential to a more effective management of global change. However, it is poorly understood how different types and intensities of extreme events create imprints in the past of ecosystems and how such legacies shape the direction of trajectories into the future. Experimental studies that employ replication and environmental control across generations are essential to gain insight in what drives alternative outcomes^4-5^. Without such technical insight humanity will not be able to repair already induced legacies before they give rise to degraded ecological states. We assembled relatively simple ecological communities in microcosms to develop theory in this area. These communities were initially comprised of three algal species. Thirty of these systems were then used to investigate how heatwaves and herbivory change the course of history by means of legacy construction. All 30 replicates first experienced a 7-day shared history under identical conditions, at a temperature of 15 °C. For the 11-day disturbance period that followed, replicates were equally divided over a control at 15 °C, heatwave treatments at 29 °C or 39 °C and herbivory treatments involving either a small-bodied or a large-bodied cladoceran herbivore species at 15 °C. At the end of the disturbance period all herbivores were removed, and the temperature was at 15 °C again in all 30 replicates until the end of the experiment. Into each post-disturbance community a herbivorous rotifer species, hereafter called the consumer, was introduced and its dynamics were observed for a period of 42 days. This design allows us to test whether past conditions shape the direction of consumer trajectories into the future, under laboratory conditions that are identical in all 30 replicates, for a period of six consumer generation times^6^.

During the pre-disturbance period all producers grew from low equal initial densities to high abundances. This resulted in an identical dominance ranking of the three producer species in all 30 replicates on day 8 of the experiment (1^st^ *Chlorella vulgaris*, 2^nd^ *Scenedesmus obliquus*, 3^rd^ *Monoraphidium minutum*, Figure 1f-h). Then an 11-day disturbance period followed, and in the subsequent 42-day post-disturbance period we investigated whether consumer dynamics differed between the control and each of the four treatments.

**Figure 1.**
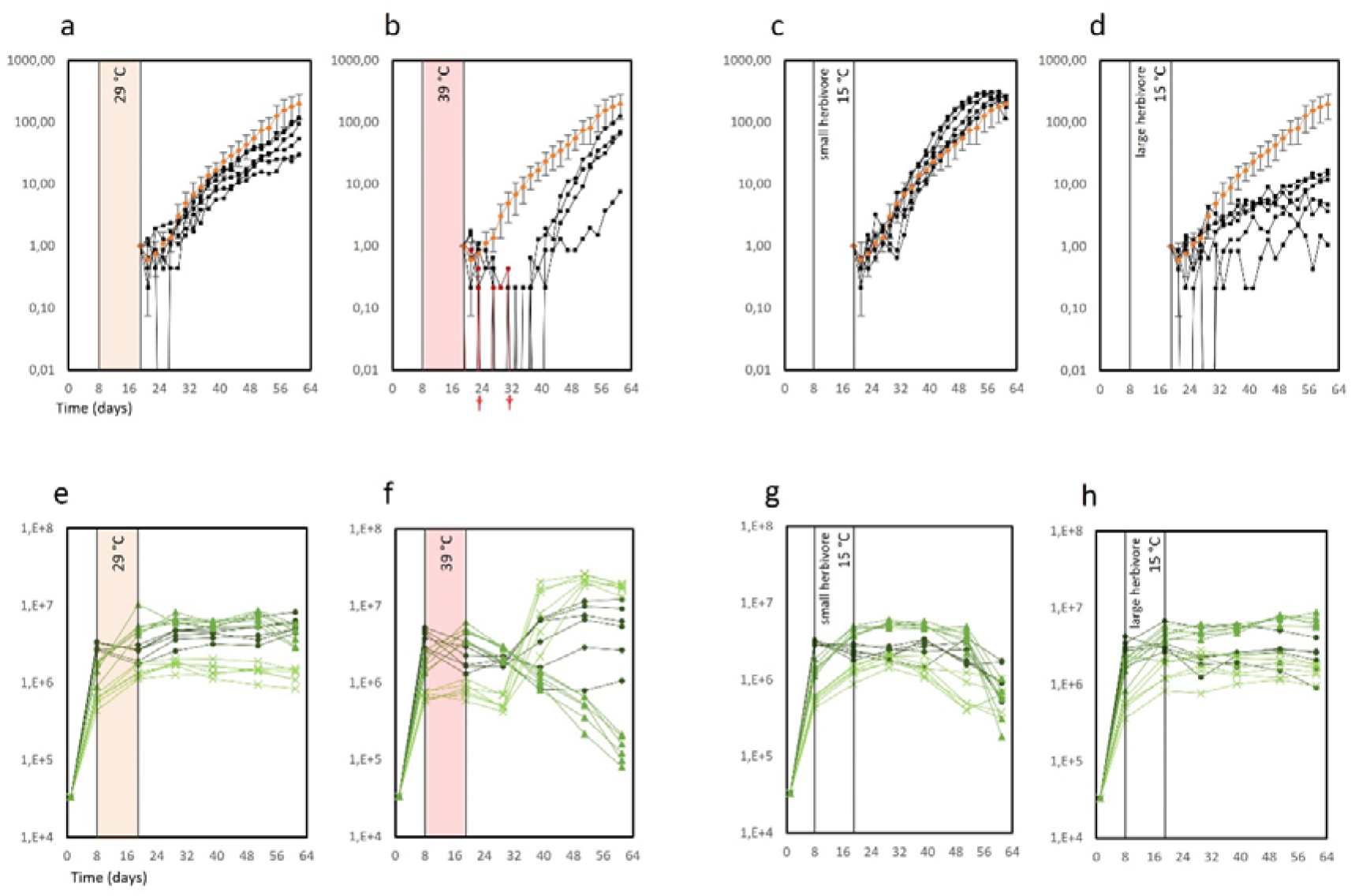
Post-disturbance dynamics of replicated consumer populations (individuals / ml). Each consumer population was introduced in a primary producer community that was previously disturbed by one of four treatments: (a) 29°C heatwave, (b) 39°C heatwave, (c) herbivory by a small-bodied cladoceran, (d) herbivory by a large-bodied cladoceran. Panels e-h show corresponding dynamics in the primary producer community (individuals / ml). Symbols legend for the primary producers, in green: x *Monoraphidium minutum*, • *Chlorella vulgaris*, and ▲*Scenedesmus obliquus*. Average consumer densities in the control are shown as orange ◊ with error bars denoting standard deviations (a-d). Trajectories leading to extinction are shown in red and marked with a cross. Consumer densities below the detection limit, represented by 0 individuals in a 4.67 ml sample (see Methods), are indicated by lines crossing the abscissa-axis.

In the control, the consumer population grew exponentially, without any extinctions, in all six replicates. A low density, below detection limit, occurred on only one single day, just after inoculation, in one of the six replicates.

A similar pattern was observed in communities preconditioned by the 29 °C heatwave. The consumer population grew exponentially, without any extinctions, in all six replicates (Figure 1a). The population was only below detection limit on a single day, in one of the six replicates (Figure 1a). Near the end of the experiment the trajectories seemed to stay a bit below the control (Figure 1a). However, there was no significant difference between this treatment and the control on any sampling day, so we conclude that the relatively mild 29 °C heatwave did not result in a legacy effect.

In contrast, in communities preconditioned by the 39 °C heatwave the consumer population crashed to a near-zero density in all six replicates (Figure 1b). This led to the consumer’s extinction in two of the six replicates (Figure 1b). Examination of the entire volume of these two microcosms at the end of the experiment confirmed that the consumer had gone extinct. One of the four remaining populations came back up but failed to grow in the direction of the control (Figure 1b). The other three did show such growth, but the difference in consumer densities between this treatment and the control was still significant at the end of the experiment. In fact, the difference in consumer density between the 39 °C history treatment and the control was significant for all sampling days from day 27 to day 61, so we conclude that extreme heat was a Legacy Constructing Factor (LCF). The extreme heat legacy took effect after a time lag of one consumer generation time (ca 7 days) and remained significant during a period of five generation times (until the end of the experiment).

Following grazing by the small-bodied herbivore, the consumer population showed exponential growth in all six replicates and then decreased somewhat, fully converging with the control at the end of the experiment (Figure 1c). There were no extinctions and consumer density stayed above the detection limit in all six replicates (Figure 1c). No significant difference in density between this past herbivory treatment and the control was detected on any of the sampling days. We hence conclude that grazing by the small-bodied herbivore did not lead to a legacy effect.

In contrast, in communities preconditioned by grazing by the large-bodied herbivore the consumer population failed to grow to a high density in all six replicates (Figure 1d). Consumer population trajectories clearly took a different direction than those of the control (Figure 1d). The consumer was shortly undetectable in two of the six replicates, but this did not lead to extinctions (Figure 1d). The difference in consumer density between this past herbivory treatment and the control was significant for 10 of the 11 sampling days from day 35 to day 61 of the experiment. Therefore, we conclude that the large-bodied herbivore was a Legacy Constructing Species (LCS). The legacy took effect after a time lag of two consumer generation times and remained significant during a period of four generation times. The legacy effect was still present at the end of the experiment, six generation times after the disturbance period had ended.

The distinct changes that we saw in consumer trajectories were a clear amplification of more modest change at the level of primary producers. Surprisingly, even the 39 °C heatwave had only a subtle effect on the producer community. It did not cause a significant difference in density between treatment and control for two of the three producer species. Only the density of the third producer was more than halved relative to the control, at the end of the heatwave. This difference was significant (P= 0.001). We know that all three producer species are edible to the consumer as we have successfully cultured it with each of them as its sole food source. However, the producer species that was significantly reduced by extreme heat was apparently its highest quality food source. When the consumer population declined to near-zero density in the post-39 °C period, this algal species took over the producer community and became dominant in all six replicates. This could be interpreted as a consumer-mediated shift in the primary producer community. This alga kept its new role as the top-ranking producer until the end of the experiment, irrespective of whether the consumer’s population grew fast, slow, or suffered extinction (Fig 1b, f).

The much stronger effects on consumers than on producers make it likely that part of the legacy that was created involves their quality as a food source. We for instance noted that algae were a paler shade of green during part of the 39 °C heatwave. Preconditioning of the producer community by extreme heat caused consumer fecundity (measured by the number of eggs per adult) to drop to a near-zero value for a substantial period (from day 21 to day 31) in the post 39 °C environment. This is one reason why the consumer population crashed so deeply in those post-heatwave producer communities. This effect on fecundity did not occur in communities preconditioned by the large-bodied herbivore.

Our results show that legacy effects from the past loaded the dice that shaped the future of ecological trajectories in our experiment. This ‘loading of dice’ occurred through a complex causal sequence. This sequence involved deterministic and chance components, time-lags, stressor amplification and major shifts in community composition. For the case of extreme heat, the nature of this sequence was to strongly amplify the initial effect. At first, only a minor change happened in the primary producer community. However, consumers responded to this change with dramatic declines leading to stochastic extinctions. These consumer declines, in turn, then had a much larger effect on the producers than the initial heatwave itself, leading to a new state of the producer community. How legacies of heatwaves and other extremes drive such causal sequences, and how these determine the future in more complex ecosystems are questions that require more scientific attention. In our simple experiment, the legacy of extreme heat drove the chance of extinction to a value of two out of six. Although this does not directly translate to the actual vulnerability of natural and man-used ecosystems to heat, we still consider this to be a high number that merits concern. It seems difficult for humanity to prevent heatwaves and further climate change, but attempting this may ultimately be easier than having to deal with the legacy effects they induce.

## Methods

### Experimental design

We used thirty 250-ml Erlenmeyer flasks as microcosms. Each of these contained 50 ml of COMBO medium with equal initial densities of three algal primary producer species, *Monoraphidium minutum, Chlorella vulgaris*, and *Scenedesmus obliquus*. Total initial density was 10^5^ individuals / ml for the primary producer community as a whole. All thirty microcosms were exposed to an acclimation and growth period of seven days under the same baseline control conditions: a constant temperature of 15 °C, gentle shaking at 60 rpm and constant LED daylight in Multitron incubators. Following this period that allowed primary producer community development, a disturbance phase of 11 days followed, from day 8 to day 19. This phase involved four treatments and a control, the latter continuing the above baseline conditions. Treatments included (i) a mild heatwave (29 °C), (ii) an extreme heatwave (39 °C), (iii) herbivory by a small-bodied cladoceran (*Ceriodaphnia dubia*), and (iv) herbivory by a large-bodied cladoceran (*Daphnia magna*). The start and end of each heatwave were implemented in steps. Temperature increase and decrease were implemented in steps of 3.5 °C per 12 hours in the 29 °C treatment. Stepsize was 6°C per 12 hours in the 39 °C treatment. On the morning of day 19, the cladoceran herbivores were removed and temperature was at 15 °C again, in all replicates. A consumer, the rotifer *Euchlanis dilatata* was then introduced at a density of 1 individual / ml (i.e. 50 per microcosm). These consumers only experienced control conditions throughout the entire post-disturbance period (day 19 to day 61), and, within the four treatments, possibly the legacy of previous exposure of the primary producer community to the different kinds of disturbance.

### Sampling & Counting

Dynamics of the consumer and all producer populations were monitored by sampling every other day. Sampling involved taking 10% of each microcosm’s volume and replacing it with fresh medium. To each 5 ml sample 2,5 ml of Lugol’s solution was added to conserve the organisms. Of the resulting 7.5 ml, 0.5 ml was used to count primary producers. The remaining 7 ml were used to count consumers. Consumer counts were thus based on 4.67 ml of each original 5 ml sample. This means that density was below our detection limit if there was less than 1 individual per 4.67 ml of medium.

## Data analysis

We first used Kruskal-Wallis tests to test for an overall difference among the five groups, (four treatments and a control), on different sampling days. In case of a significant overall difference, each of the four treatments was compared with the control using a Dunn post-hoc test that had its p-value adjusted by the Benjamini-Hochberg FDR method. This procedure statistically corrected for each of the four treatments being compared with a single control.

## Acknowledgement

We gratefully acknowledge funding to MV by the Deutsche Forschungsgemeinschaft (DFG, German Research Foundation) – SFB 1439/1 2021 – 426547801.

## Notes

### Competing Interest Statement

The authors have declared no competing interest.

